# Precocious sperm exchange in the simultaneously hermaphroditic nudibranch, *Berghia stephanieae*

**DOI:** 10.1101/2022.03.02.482740

**Authors:** Neville F. Taraporevala, Maryna P. Lesoway, Jessica A. Goodheart, Deirdre C. Lyons

## Abstract

Sexual systems vary greatly across molluscs. This diversity includes simultaneous hermaphroditism, with both sexes functional at the same time. Most nudibranch molluscs are thought to be simultaneous hermaphrodites, but detailed studies of reproductive development and timing remain rare as most species cannot be cultured in the lab. The aeolid nudibranch, *Berghia stephanieae*, is one such species that can be cultured through multiple generations on the benchtop. We studied *B. stephanieae* reproductive timing to establish when animals first exchange sperm and how long sperm can be stored. We isolated age- and size-matched individuals at sequential timepoints to learn how early individuals exchange sperm. Individuals isolated at 13 weeks post laying (wpl) can produce fertilized eggs. This is 6 weeks before animals first lay egg masses, indicating that sperm exchange occurs well before individuals are capable of laying eggs. Our results indicate that male gonads become functional for animals between 6 mm (~9 wpl) and 9 mm (~15 wpl) in length. That is much smaller (and sooner) than the size (and age) of individuals at first laying (12-19 mm; ~19 wpl), indicating that male and female functions do not develop simultaneously. We also tracked the number of fertilized eggs in each egg mass, which remained steady for the first 10-15 egg masses, followed by a decline to near-to-no fertilization. This large, novel dataset provides insights into precise timing of the onset of functionality of the male and female reproductive systems in *B. stephanieae*. These data contribute to a broader understanding of reproductive development and the potential for understanding evolution of diverse sexual systems in molluscs.

## Introduction

Sexual systems–the patterns of sex allocation within a species–are diverse across animal lineages. While most animals have separate sexes (gonochorism, dioecy), hermaphroditism (monoecy) occurs in 5% of all described species, increasing to 30% when insects are excluded (Jarne and Auld 2006). Most major metazoan lineages have evolved hermaphroditism, which includes two major categories: simultaneous and sequential hermaphroditism, with female and male function occurring either at the same time, or at different times, respectively. Current hypotheses suggest that reproductive plasticity plays an important role in the evolution of sexual systems (Leonard 2013, 2018).

Although definitions of hermaphroditism imply sharp distinctions between sequential and simultaneous hermaphroditism, the underlying biology is not so clear-cut (Heller 1993; Collin 2013; Leonard 2018). Apportioning of reproductive output between male and female functions in simultaneous hermaphrodites often changes in response to environmental inputs, including social and physical environment (Tomiyama 1996; Yusa 1996; Lorenzi et al. 2006; Baeza 2007; Janicke and Schärer 2009; Ramm et al. 2019). However, relatively little empirical work has addressed the biology of simultaneous hermaphrodites in comparison to studies of separate sexes, particularly in groups where transitions among different sexual systems are more common. In order to understand how transitions between sexual systems might occur, more detailed anatomical and reproductive data are needed (Nakadera and Koene 2013). In particular, longitudinal reproduction data in simultaneous hermaphrodites, including how and when reproductive resources are allocated, will be important to understanding transitions between sexual systems.

Molluscs are an excellent group in which to study reproductive diversification and the mechanisms underlying transitions between sexual systems. Molluscan sexual systems are highly labile, and include multiple origins of simultaneous and sequential hermaphroditism, with the greatest diversity of sexual systems in molluscs found in gastropods and bivalves (Collin 2013, 2018; Lesoway and Henry 2019). Within gastropods, separate sexes are thought to be ancestral, and both simultaneous and sequential hermaphroditism have evolved repeatedly (Collin 2013). For example, all members of the family Calyptraeidae are sequential hermaphrodites (Collin 2013, 2018; Lesoway and Henry 2019), and the subclass Heterobranchia, which includes both land snails (pulmonates) and sea slugs (*e.g.,* nudibranchs), is considered (primarily) simultaneously hermaphroditic (Heller 1993; Jarne and Auld 2006).

Within Heterobranchia, nudibranchs have proven to be useful for studies of development (Bonar and Hadfield 1974; Carroll and Kempf 1990; Kempf et al. 1997), animal behavior and neurobiology (Willows 1971; Kriegstein et al. 1974; Chase 2002; Katz and Quinlan 2019), nematocyst sequestration (Goodheart and Bely 2017; Goodheart et al. 2018, 2022), and reproduction (Rutowski 1983; Rivest 1984; Hall and Todd 1986; Angeloni et al. 2003; Sekizawa et al. 2019). Nudibranchs are typically described as simultaneous hermaphrodites with concurrently functioning male and female sex organs and reciprocal sperm exchange between conspecifics during mating (Hall and Todd 1986; Karlsson and Haase 2002; Angeloni et al. 2003; Sekizawa et al. 2019). However, descriptions of reproduction in most hermaphroditic species are often based on limited sampling of a few adult individuals (Collin 2013), and several sources describe nudibranchs as having a more complex sexual system than simply simultaneous hermaphroditism (Tyrell Smith and Carefoot 1967; Harris 1975; Todd et al. 1997; Wägele and Willan 2000). Indeed, the reproductive diversity of nudibranchs appears to be something of an open secret in the nudibranch literature (Hadfield and Switzer-Dunlap 1984; Heller 1993; Wägele and Willan 2000), but supporting data remains limited to a single species, *Phestilla sibogae* (Todd et al. 1997). We are not aware of other studies that address the timing of gonadal development in nudibranchs, leaving open many questions about how and when reproductive resources are allocated in members of this group. However, our understanding of nudibranch biology is limited by the challenges of culturing nudibranchs through the complete life cycle in the lab. The aeolid nudibranch, *Berghia stephanieae* (Fig. 1A), can be kept easily in the lab, and presents a convenient research organism for studying nudibranch reproduction and development (Carroll and Kempf 1990; Kristof and Klussmann-Kolb 2010; Dionísio et al. 2013).

**Figure 1.**
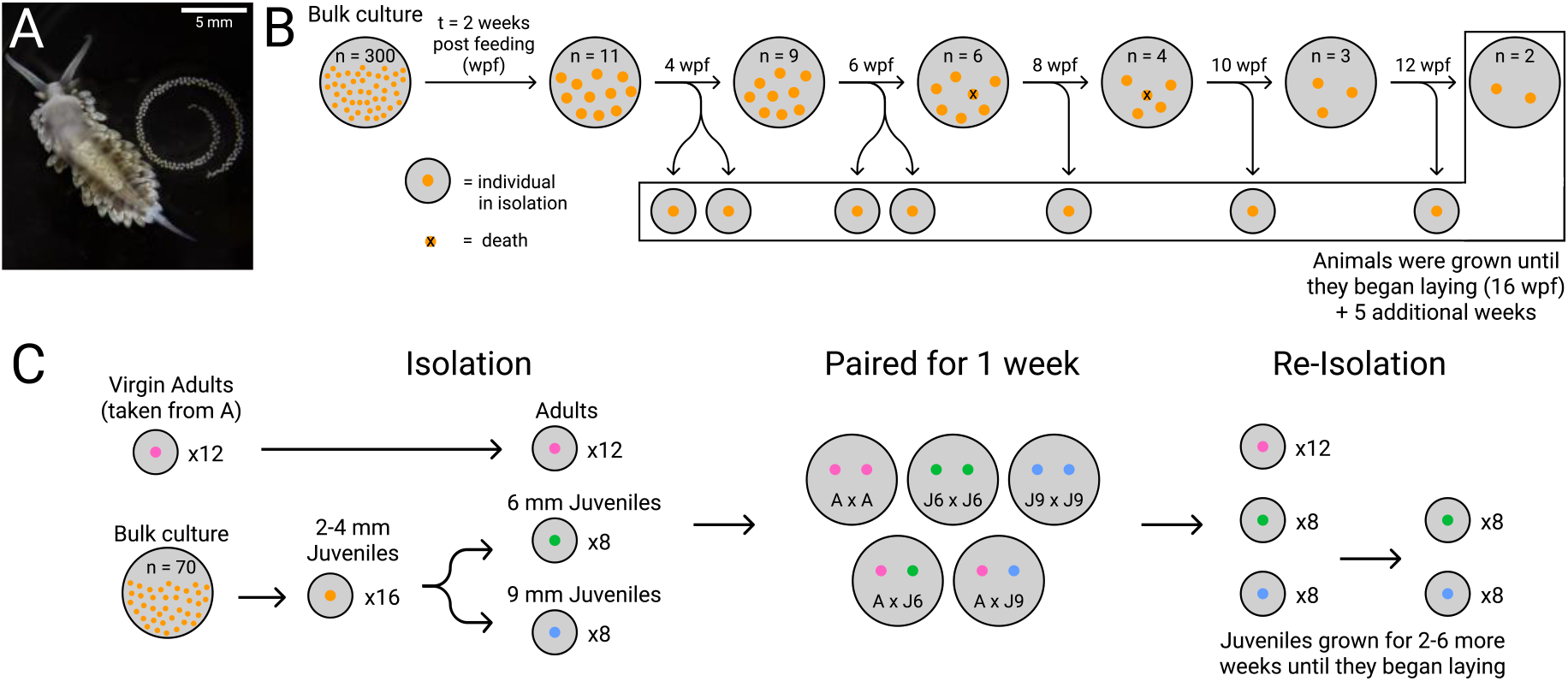
Experimental Design for Reproductive Isolation Experiments. **A.** Adult *B. stephanieae* in ventral view, with egg mass. Scale = 5mm. **B.** Approximate age and size matched juveniles from the initial bulk culture were separated into six groups of 11 animals (n = 66 total individuals) which were reared for a total of 23 weeks. Juveniles (n = 40) were then sequentially isolated over a period of 8 weeks. On average, 2-3 individuals died from each group over the isolation period. **C.** New individuals and virgin adults from the first experiment (A) were grown in isolation, paired for one week, then re-isolated. The number of fertilized and unfertilized eggs were collected for egg masses produced by isolated animals. Colored disks indicate single individuals, x indicates dead individuals.

The availability of long-term cultures of *B. stephanieae* allows us to ask questions about when sperm is produced and exchanged, how sperm is allocated, and allows us to analyze changes in reproductive output over time. In this manuscript, we ask: (1) how early are individuals able to exchange sperm and can individuals exchange sperm at the same time that they produce fertilized eggs; (2) how long isolated individuals are able to retain sperm; and (3) how does sperm quality change over time? To determine the earliest stage at which these animals exchange sperm, we separated individuals at sequential timepoints. To confirm our results, we also performed reciprocal mating experiments on isolated individuals of different sizes, in addition to confirming our observations with histological sectioning of reproductive tissues across multiple reproductive timepoints. Together, our results show that individuals can produce and exchange sperm prior to producing eggs, documenting an additional species of nudibranch with individuals that function first as males, then as simultaneous hermaphrodites. In addition, we report longer sperm storage times than previously reported for other nudibranchs, and document changes in fertilization rates over time.

## Methods

### Adult cultures and juvenile collection

Adults of *Berghia stephanieae* Valdés 2005 (Fig. 1A) were originally acquired from Reeftown (https://Reeftown.com), and then maintained in continuous culture in our laboratory. Briefly, animals were kept in large finger bowls in artificial seawater at a salinity of 1.024 sg (Instant Ocean, Spectrum Brands, Blacksburg, VA) at room temperature (~20°C) and allowed to mate freely. Groups and individuals were fed enough *Exaiptasia diaphana* (Carolina Biological Supply, Burlington, NC) to develop properly (see Suppl. File 1), and in order to prevent cannibalism.

### Age-based isolation

In our first experiment, juveniles of known age were used for controlled matings. First, egg masses were collected within a five-day window and allowed to hatch and metamorphose (~ 2-3 weeks post laying, wpl) into juveniles in a large fingerbowl. A group of ~300 of these recently metamorphosed juveniles (~ 3 wpl) were then fed together prior to experimental isolation. In these cultures, development is asynchronous, as larger animals consuming most of the food can limit growth of smaller individuals (Monteiro et al. 2020). To minimize such asynchronous development, after 2 weeks of feeding (2 weeks post feeding, wpf), the 66 largest individuals (~1-2 mm long) were removed from the bulk culture in the finger bowls and raised in 6-well plates (Corning, #3736), grouped by size into 6 sub-groups of 11 individuals each (Fig. 1B). At 4 weeks post feeding (wpf, ~7 wpl), 2 individuals from each sub-group (12 total) were isolated to their own separate wells to prevent access to additional mating events. Subsequently, 12, 6, 6, and 4 individuals were likewise each separated to individual wells at 6, 8, 10, and 12 wpf, respectively. The remaining 4 individuals were kept in 2 mating pairs that continued to have mating access for the duration of the experiment.

All individuals, regardless of mating status, start producing egg masses at 16 weeks post feeding (Fig. 1B). Isolated and grouped animals were maintained for an additional 5 weeks (21 wpf) to give them sufficient time to lay several egg masses (~16 wpf; ~19 wpl). When individuals began to lay, egg masses from each animal were collected daily. Egg masses from animals isolated at 12 weeks post feeding were continuously collected until egg mass production ceased, after 16 weeks in total. Over the course of the experiment, a third of the animals cultured (22 individuals) died, leaving 44 animals for isolation experiments (Fig. 1B). Individuals that were isolated but died (n=7) before laying any egg masses and were not included in our analyses.

Egg masses were removed from plates with forceps and separated into 12-well plates (ThermoScientific, #150200) in ~3 mL of 0.22μm filtered artificial seawater for 1-2 weeks and allowed to develop. Previous experiments in the lab confirmed these as optimal conditions for development, with typically low mortality in developing embryos (> 80% metamorphosis). Embryos were then examined and categorized into unfertilized eggs, or fertilized embryos, based on their morphology. Embryos that showed stereotypical cleavage and developed into typical veligers (Fig. 2E) were categorized as fertilized. All others including non-cleaving- and chaotic-cleaving-eggs (Fig. 2B) were categorized as unfertilized. In egg masses containing fertilized eggs, all eggs, unfertilized and fertilized, were counted (Fig. 3). Egg masses without fertilized eggs were scored as being unfertilized but the number of eggs was not counted.

**Figure 2.**
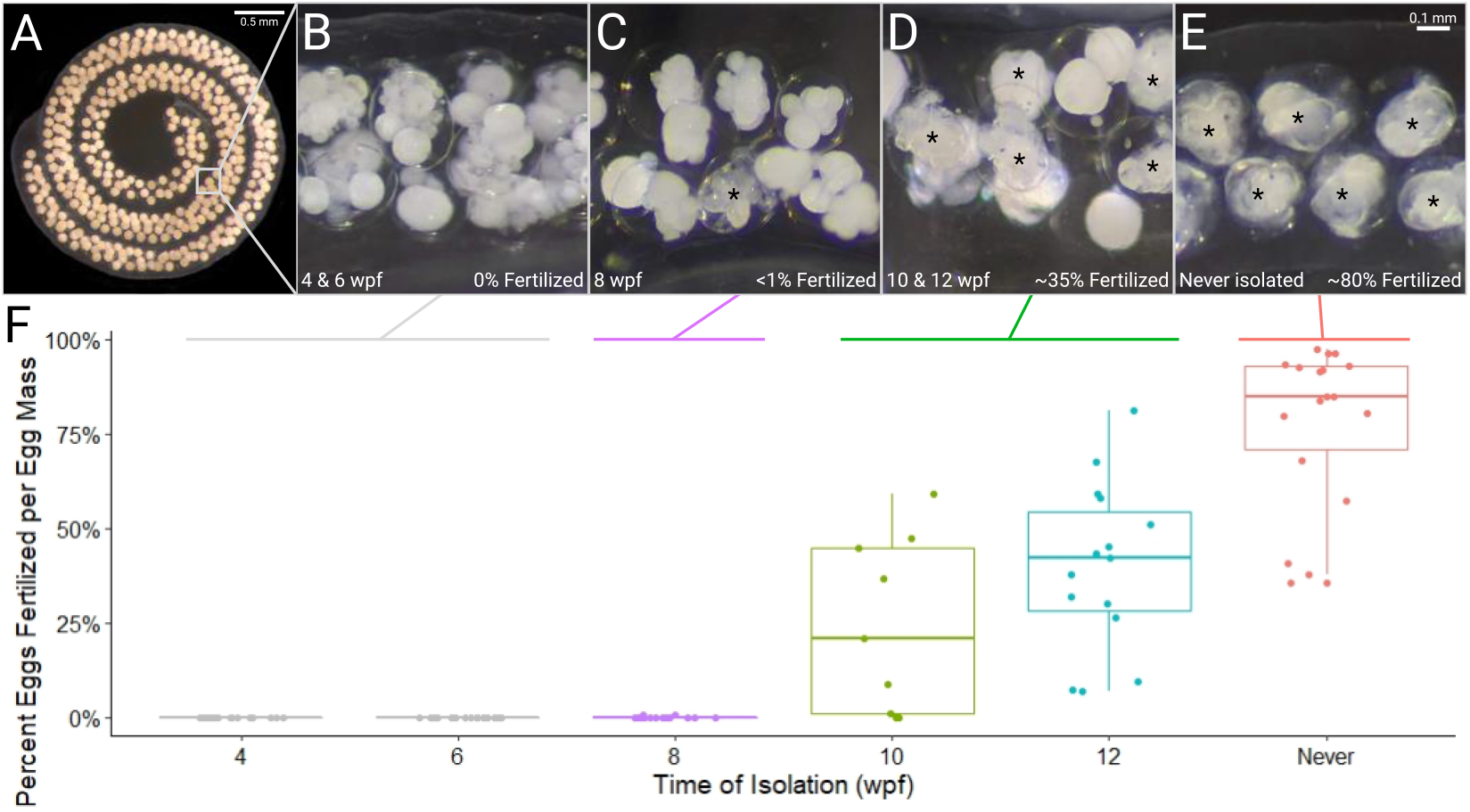
Later isolation increases fertilization rates. **A.** Coiled egg mass of *B. stephanieae* containing multiple encapsulated embryos. **B.** Unfertilized eggs from an egg mass laid by an animal isolated at 6 weeks post feeding (wpf). **C.** One fertilized egg among unfertilized eggs from an egg mass laid by an animal isolated at 8 wpf. **D.** Some fertilized eggs in an egg mass laid by an animal isolated at 12 wpf. **E.** Most eggs are fertilized in an egg mass laid by an animal that was never isolated. (Asterisk - fertilized embryo). **F.** Fertilization rate for egg masses produced by age-isolated animals, grouped by time of isolation.

**Figure 3.**
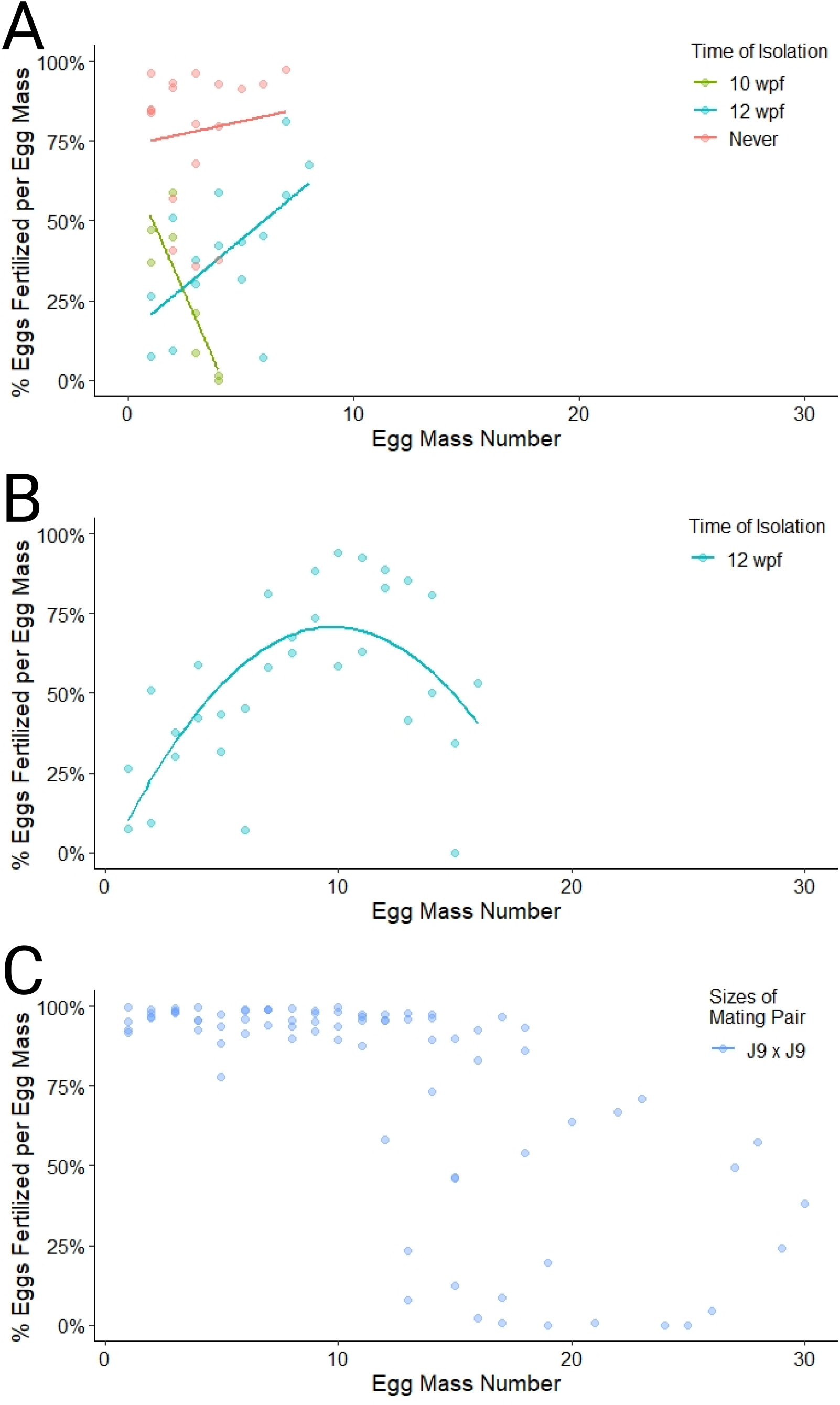
Fertilization rate changes over time in both time- and size-isolated individuals. **A.** Fertilization rate for age-isolated animals over the 5 week experimental period. Egg masses are grouped by the time individuals were isolated **B.** Fertilization rate of egg masses laid by two individuals isolated at 12 weeks post feeding (wpf) over a 12 week period (continued from A). **C.** Fertilization rate of egg masses produced by large juveniles mated with large juveniles (9mm, LJxLJ) in the size selection experiment.

### Size-based isolation

Development in *B. stephanieae* is highly asynchronous, and varies depending on environmental conditions (i.e. food availability, temperature, inter-individual feeding competition), making estimations of age in bulk culture challenging. As demonstrated above, tracking precise age is time consuming and labor intensive. We therefore performed a second experiment isolating animals by size, which can be assessed easily (Fig. 1C). Based on the results of our first experiment, we estimated size classes of animals able to produce and exchange sperm. We determined three categories of animals: small juveniles (5-6 mm, equivalent to animals from age-based experiment ~6 wpf/~9 wpl, Fig. 4A), large juveniles (over 9 mm, ~12 wpf/~15 wpl, Fig. 4B), and adults (>12 mm, >16 wpf/>19 wpl, Fig. 4C). We expected small juveniles not to be able to exchange sperm, and large juveniles and adults to exchange sperm. Adults (n = 12) were taken from our previous experiment. These adult animals, isolated at 4 and 6 wpf, were presumed to have never received sperm since they laid only unfertilized egg masses, and were over 15 mm in length. The juveniles (2-4 mm in length, n = 16) were selected from a new bulk culture and put into isolation until they reached the appropriate sizes for our experiment (5-6 mm and >9 mm; Fig. 1C).

**Figure 4.**
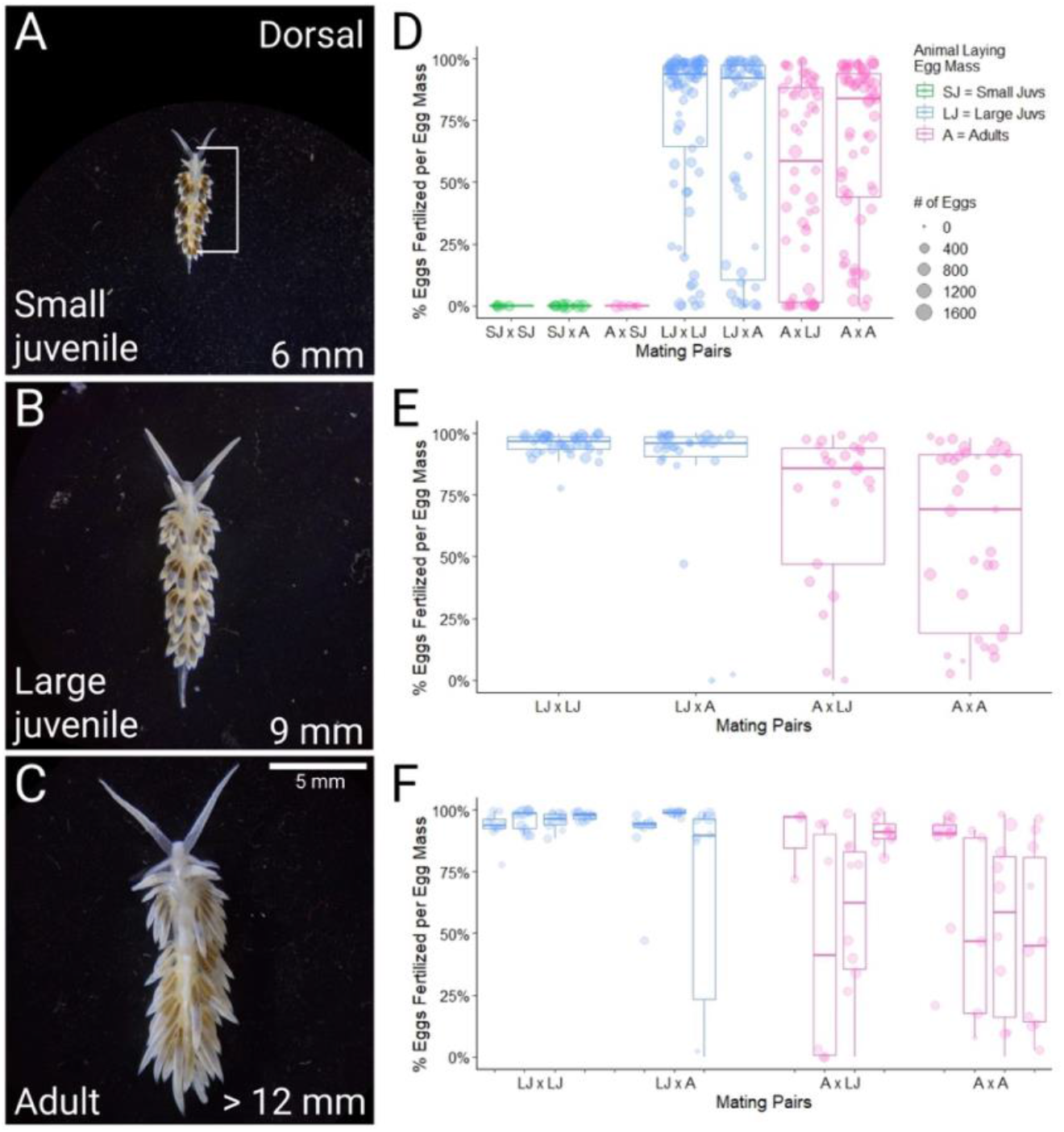
Fertilization By Size Category. **A-C.** Dorsal view of *B. stephanieae* at 3 lengths measured from mouth to end of cerata. **A.** Small juvenile (6 mm). **B.** Large juvenile (9 mm). **C.** Adult (~12-20 mm). Scale bar = 5 mm. **D.** Fertilization rate of all egg masses laid by all animals in the size-based experiment. The first individual in the cross is the animal that laid the egg mass and second is the available mate. **E.** The first 10 egg masses laid in sequence by adult and large juveniles, categorized by the animal they were paired with. **F.** Individual variation across the first 10 egg masses laid in sequence by each individual, organized by cross type.

Animals were measured from photographs taken while fully extended during active crawling, from the anterior-most part of the head to the furthest posterior cerata (Fig. 3A). Photographs of each animal were taken using a dissecting scope (Zeiss Stemi 305) and a Google Pixel 3A smartphone. Five separate images were taken of each animal, and the length of the animal was measured in each of these images using Fiji (Schindelin et al. 2012).

In total, 14 pairs were mated, including 2 pairs each of virgin adults, small juveniles, and large juveniles, and 4 pairs each of virgin adults with either small or large juveniles (Fig. 1C). Paired animals were allowed to mate for 7 days and were not fed, to minimize growth. When two animals of the same category were paired, a small clump of cerata (finger-like projections of the dorsum) was removed from one animal to differentiate the two individuals. The animals were then re-isolated and fed one-twelfth of a medium-sized *E. diaphana* every other day for the remainder of the experiment (2-4 months). Adults (>7 months post laying) began laying egg masses immediately, while large and small juveniles (>2 months post laying) started laying egg masses ~2 and ~6 weeks after re-isolation, respectively (Fig. 1C). Egg masses were collected following the same procedure as described above. Additionally, one juvenile and two adults were kept in isolation for the duration of the experiment as controls.

### Statistical analyses

All statistical analyses were performed using R (v.4.04). The relationship between the fertilization rate of egg masses (dependent variable) and the order of egg masses laid (independent variable) was analyzed with a linear regression (lm function in stats v. 4.0.3; Fig. 3A) for each time-based isolation treatment (10 wpf, 12 wpf, and never isolated individuals). For the size based experiments, differences in fertilization rates among mating categories (Fig. 4A) were analyzed using an ANOVA (car v. 3.0-10; Fig. 4A), followed by Tukey HSD tests to assess which specific mating category means were significantly different. Individual variation within mating categories was then analyzed with an ANOVA (car v. 3.0-10; Fig. 4B-C). Graphs were prepared using ggplot2 (v. 3.3.5). Figures were prepared using Figma (https://figma.com). All relevant scripts and raw data files are available on GitHub (<link to be added on manuscript acceptance>).

### Histology and Imaging

Animals matching the previously determined size categories (6 mm, 9 mm, >12 mm) were measured and selected from bulk cultures, as described above. Animals were relaxed in a 1 part 7.5% MgCl_2_ to 2 parts Artificial Sea Water (ASW) solution until movement ceased, and then fixed in 4% paraformaldehyde in filtered sea water solution overnight at 4°C. A post-fixation stain was then performed using Ponceau S (0.1% Ponceau S and 1.0% glacial acetic acid) for 2 hours, followed by a diH_2_O rinse. The tissues were then dehydrated: (1) 50% ethanol (EtOH) for 15 min, (2) 60% EtOH for 15 min, and (3) 70% EtOH for 15 minutes. Tissues were then stored in 70% EtOH (at 4°C) or immediately prepared for embedding. In preparation for embedding, we subjected tissues to a further dehydration series: (1) 80% EtOH for 15 min, (2) 95% EtOH for 15 min, (3) 100% EtOH for 15 minutes (x2). Samples were embedded with Spurr's Low Viscosity Embedding Media Kit (Electron Microscopy Sciences #14300), following the “Firm” formulation provided by the manufacturer, and cured at 60°C overnight. Sections 3μm thick were cut using glass knives on a microtome, and sections were mounted and counterstained with Azure A. Sections were imaged using a Zeiss AxioM2 fluorescence microscope with an attached digital camera and ZEN software (v2.3). Image stacks were combined in Helicon Focus (v7.7.5), adjusted for brightness and contrast using Fiji (v1.53f51, Schindelin et al. 2012). Image plates were prepared using the FigureJ plugin (Mutterer and Zinck 2013), https://imagej.net/plugins/figurej) and Adobe Illustrator (v26.0.3).

## Results

### Age-based Isolation: Later isolation increases fertilization rates

Successive isolation of carefully age-and size-matched *Berghia stephanieae* juveniles indicates the approximate age that individuals are able to exchange sperm (Fig. 2). The earliest isolated individuals (4 and 6 weeks post feeding; wpf) laid only unfertilized eggs (n = 42 egg masses, Fig. 2A). Of the 15 egg masses (>1000 eggs total) laid by animals isolated at 8 weeks post feeding, only a single egg was fertilized (Fig. 2B). Juveniles isolated at 10 and 12 wpf consistently produced fertilized eggs (Fig. 2C, E), laying 10 and 15 egg masses, respectively, with all but 2 containing fertilized eggs. Animals that were never isolated laid 17 egg masses, all of which contained fertilized eggs.

Fertilization rates varied among treatments, and animals with continuous access to sperm had significantly higher fertilization rates compared to individuals isolated at 10 and 12 wpf (p ≪0.001 for each). For animals that were never isolated, average fertilization was high (78.0 ± 21.1%, Fig. 2E), while animals isolated at 10 and 12 wpf laid egg masses with low average fertilization rates (24.3 ± 23.2% and 39.8 ± 22.0%, Fig. 2E). The long duration of these experiments also allowed us to observe changes in fertilization rates over time. For the animals that were never isolated, the proportion of eggs fertilized in each egg mass did not significantly change over time (r^2^ = 0, p = 0.62, Fig. 3A). Two animals isolated at 10 wpf laid multiple fertilized egg masses. In these animals, fertilization declined significantly (r^2^ = 0.63, p = 0.008, Fig. 3A). For the animals isolated at 12 wpf, fertilization rates measured during the initial 5-week experimental period increased (r^2^ = 0.31, p =0.017, Fig. 3A). We therefore continued to measure fertilization rates in these animals for a further 7 weeks. Over the course of 13 weeks, fertilization rates of egg masses laid by animals isolated at 12 wpf best fit a quadratic equation (r^2^ = 0.41, p < 0.001, Fig. 3B), with the highest fertilization rate around the 10th egg mass laid (Fig. 3B).

### Size-based Isolation: large juveniles are able to exchange sperm with adults and each other

Asynchrony in juvenile development in *B. stephanieae* makes estimating age difficult, so in order to make our results more broadly applicable to our batch cultures, we turned to size as a proxy for reproductive stage. Guided by our previous experimental results, we defined three size classes corresponding to immature juveniles, animals capable of exchanging sperm and and mature animals capable of producing fertilized eggs: small juveniles (6mm long), large juveniles (9 mm long) and adults (>12 mm long) (Fig. 4A-C). Adults that were paired either with large juveniles (>9 mm) or with other adults produced fertilized egg masses almost immediately (Fig. 2C), while large juveniles paired with both adults and other large juveniles produced egg masses after a few weeks of additional growth (Fig. 2C). When animals were mated based on size, sperm exchange occurred in animals 9 mm and larger. Small juvenile pairings (<6 mm at the time of mating; SJxSJ) produced almost no fertilized egg masses (only a single egg was fertilized out of 28 egg masses, (Fig. 4A), while large juvenile pairings (9 mm; LJxLJ) and adult pairings (>12 mm; AxA) produced primarily fertilized egg masses (Fig. 4A,). Small juveniles paired with adults produced almost no fertilized eggs (Fig. 4A; SJxA), with the exception of a single fertilized egg produced by one animal in the second of 3 otherwise unfertilized egg masses. Similarly, adults that were paired with small juveniles did not produce any fertilized eggs (Fig. 4A; AxSJ). Overall, both adults and large juveniles were able to exchange sperm while small juveniles were not.

Fertilization rates of the egg masses produced when animals were isolated based on size significantly correlated with the size of the animal laying the egg mass. Because the fertilization rate dropped significantly starting around the 10th to 15th egg mass laid (Fig. 3C), presumably due to animals “running out of sperm”, we considered only the first ten egg masses produced by each animal to test for differences in fertilization rates between adults and large juveniles. Fertilization rates of egg masses laid by adults mating with other adults were the lowest (not including small juveniles) and most variable (59.1% ± 35.5%, Fig. 4B), followed by adults mating with large juveniles (69.6% ± 34.1%), then large juveniles mating with adults (84.6% ± 30.0%), then large juveniles mating with other large juveniles (95.5% ± 4.3%, Fig. 4B). Large juveniles laid egg masses with higher fertilization rates than adults regardless of whether the mate was an adult (p = 0.002, Fig. 4B) or a large juvenile (p = 0.001, Fig. 4B). However, the age of the mate did not appear to change fertilization rates. Regardless of whether the mate was an adult or a large juvenile, fertilization rates were similar for egg masses laid by adults (p = 0.44, Fig. 4B) and large juveniles (p = 0.38, Fig. 4B). As noted, the percent of eggs that are fertilized in each egg mass remains steady for the first ~10 egg masses on average followed by a decline to near-to-no fertilization (Fig. 3C). While fertilization rate varied based on treatment, in all cases, fertilization eventually declined to zero.

Within groups, individuals were fairly consistent in egg mass fertilization (Fig. 4C). Among the first 10 egg masses laid, adults (n = 8) had consistently high variation and a Tukey test showed that no individual was significantly different from another (p > 0.05 for all, Fig. 4C). Large juveniles (n = 7) generally produced egg masses with consistently high fertilization rates, and a Tukey test showed only one individual was significantly different from the rest (p < 0.05, n = 1, Fig. 4C). This indicates that there is no significant variation among individuals, and adults generally showed a decline in fertilization when compared to large juveniles.

### Histology: Gonadal Development and Sperm Localization

Mating experiments using either age- or size-selected animals (above) indicate that sperm exchange occurs prior to fertilization and egg mass laying. To confirm the presence and location of sperm in the reproductive tract and the timing of sperm and egg development, we cut histological sections of small juveniles, large juveniles and adults as determined above. Histological sections showed clear differences in gonadal maturation across the small juvenile (6 mm; Fig. 5A), large juvenile (9 mm; Fig. 5B) and adult (>12 mm; Fig. 5C) stages (Fig. 5D-I). At all stages, the gonad (or ovotestis), where sperm (*i.e.,* autosperm) and eggs are produced, contained mature sperm and sperm progenitor cells (Fig. 5G-I). The ovotestis of adults also contained mature oocytes (Fig. 5I), and signs of oocyte maturation were visible in large juveniles (Fig. 5H). Mature oocytes were not observed in the reproductive tract of the individuals sectioned. Reproductive morphology of nudibranchs is complex, with structures for storage of autosperm (ampulla, data not shown) and allosperm, sperm received from mates (seminal receptacles) (Fig. 5D-F). Identification of these structures often relies on location in the animal and orientation of sperm. Autosperm are typically oriented at random within the ampulla, while allosperm are oriented with the sperm heads directed into the epithelium of the seminal receptacle (Wägele and Willan 2000). No sperm was observed outside the ovotestis in the 6mm juveniles (Fig. 5D), while the seminal receptacles and ampullae of both large juveniles and adults contained sperm (Fig. 5E-F). Together, the presence of sperm (both autosperm and allosperm) outside the gonad in large juveniles and the delayed maturation of oogonia confirm that sperm production and exchange precede egg maturation and fertilization in *B. stephanieae*.

**Figure 5.**
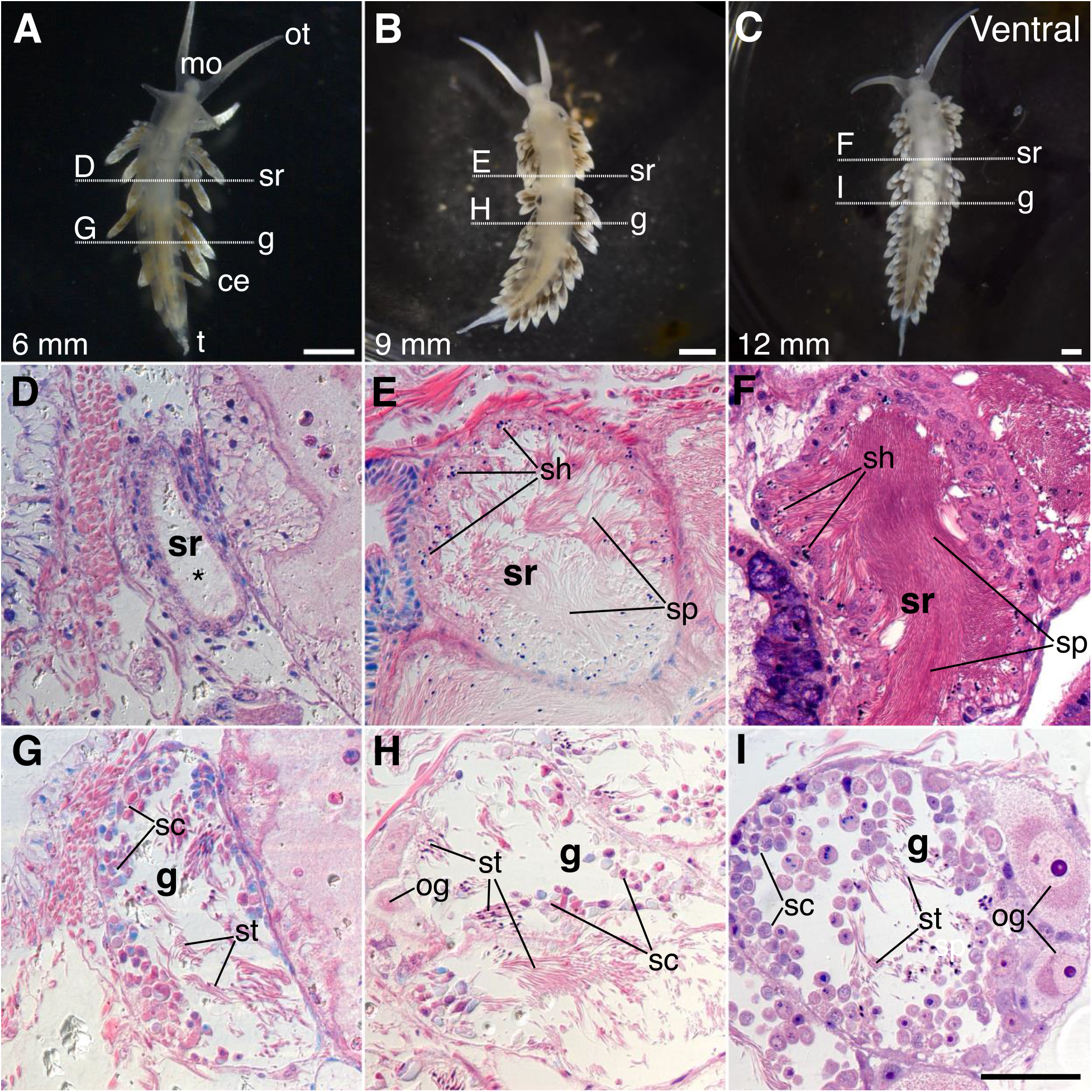
Histology of reproductive development in *Berghia stephanieae*. **A-C.** Ventral views of 6mm **(A)**, 9mm **(B)** and 12mm **(C)** juveniles of *B. stephanieae*. Horizontal lines indicate the approximate level of histological sections. **D-F.** Sections cut through the seminal receptacles show increasing amounts of stored allosperm. Sperm heads are directed towards the wall, in contrast to autosperm, which are randomly oriented within the ampulla (data not shown). No sperm was seen in the reproductive tract of the 6mm animal (**D**, asterisk) outside of the gonad (**G**). **G-I.** Histological sections cut through the developing gonads. Both spermatocytes and maturing oocytes are produced in the ovotestis. Sperm maturation **(G)** precedes oocyte maturation **(I)**. ce, cerus; g, gonad; mo, mouth; t, tail; ot, oral tentacle; sh, sperm heads; sp, sperm; st, spermatid; sc, spermatocyte; sr, seminal receptacle; oc, oocyte. Scale: A-C = 1 mm, D-I = 50 μm.

### Discussion

Our study shows that *Berghia stephanieae* are not simply simultaneous hermaphrodites, and highlights the complicated nature of sexual systems and their development. Male and female functions do not develop simultaneously in *B. stephanieae*. We find that sperm exchange in *B. stephanieae* begins between 8 and 10 weeks post feeding (wpf, 6-8 weeks before laying egg masses) and between 6 and 9 mm in length (Fig. 6). Our results also show that the ovotestis begins to produce sperm before sperm exchange starts, as spermatogenesis is evident in small (6 mm) juveniles (Fig. 5G). Our results show that sexual development is more complex than existing descriptions of simultaneous hermaphroditism in nudibranchs would suggest, and contributes detailed data about the timing and allocation of sexual resources.

**Figure 6.**
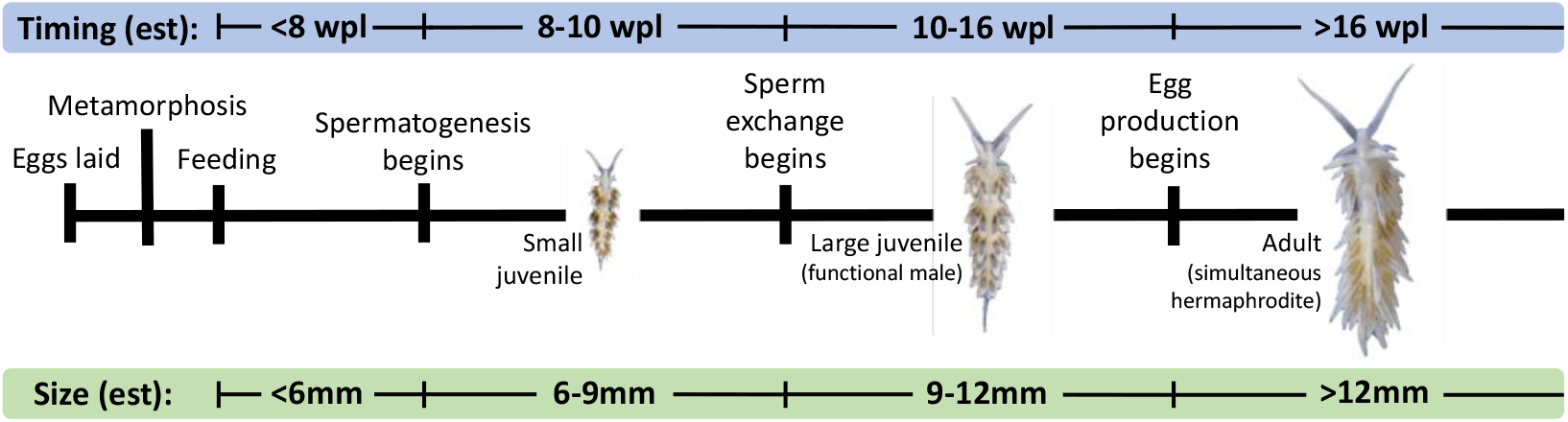
Summary of *Berghia stephanieae* reproductive development, with estimated timing (at RT, ~20°C) and approximate size ranges (mm). Small juveniles (~8-10 weeks post-laying, wpl; ~6-9mm in size) are able to produce sperm but are unable to exchange it. Large juveniles (~10-16 wpl; ~9-12mm in size) are able to both produce and exchange sperm. Adults (>16 wpl; >12mm in size) are able to produce and exchange sperm and lay fertilized egg masses. est, estimated; mm, millimeters; wpl, weeks post-laying.

### Sperm exchange occurs before egg laying

Here, we provide detailed evidence that individuals of *B. stephanieae* become functionally male before becoming simultaneous hermaphrodites. While making direct comparisons between the time-based and size-based experiments is difficult (see below), animals exchange sperm when they are at least 6 weeks younger and 3 mm smaller than when they are able to lay eggs (Fig. 6). This is consistent with previous work in the nudibranch *Phestilla sibogae*, which showed sizes of animals able to exchange sperm (5-8 mg) are 10-20x smaller than those able to lay eggs (90-480 mg, Todd et al. 1997), and in sacoglossans (another group of heterobranch gastropods), which show a similar pattern of reproductive development. Juveniles of the sacoglossan *Alderia modesta* also exchange sperm before being able to lay eggs (2 vs. 10 days post metamorphosis) at at half the size (0.5 mm vs. 1.2 mm long, Angeloni et al. 2003), and in the sacoglossan *Elysia maoria*, male gonads appear in animals only 5 mm long while female gonads begin to appear in animals 10 mm long (Reid 1964). Becoming functionally male before egg mass production begins appears to be more common in heterobranch gastropods than is widely recognized.

The phenomenon of a male phase before simultaneous hermaphroditism is best categorized as simultaneous hermaphroditism with adolescent protandry, a subset of simultaneous hermaphroditism (after Leonard 2018). Elsewhere, this pattern of male first reproduction has been described as adolescent gonochorism (Heller 1993), protandrous hermaphroditism (Wägele and Willan 2000), or “intermediate between the conventional ‘simultaneous’ and ‘sequential’ categories.” (Todd et al. 1997). Protandrous hermaphroditism implies a switch from one sex to the other including loss of function of the first sex, which is not the case here (Ghiselin 1969; Leonard 2018). This is also not likely an evolutionarily intermediate stage since sequential hermaphroditism does not appear to have evolved from simultaneous hermaphroditism in other gastropods (Collin 2013). Our results highlight the importance of documenting reproduction through the full reproductive life cycle. In order to draw conclusions about the mechanisms that underlie transitions between sexual systems, we must have a fuller understanding of reproductive allocation, particularly within simultaneous hermaphrodites (Schärer 2009; Nakadera and Koene 2013).

### Fertilization rates over time

We predicted that fertilization rates would decrease uniformly, starting high and then going to zero as an individual “ran out” of sperm. For example, in the nudibranch *Hermissenda crassicornis*, percent fertilization decreased as more egg masses were produced within an individual (Rutowski 1983). Similarly, fertilization rates of singly-mated individuals of the heterobranch gastropod *Aplysia californica* started at high levels and declined as egg mass production continued (Ludwig and Walsh 2004). While this was generally the case in our study, not all of our findings showed an immediate decline. Egg masses laid by the 2 animals isolated at 12 weeks post feeding show an initial increase in fertilization before decreasing, after producing approximately 10 egg masses (Fig. 3 A-B), while large (9 mm) juveniles produced egg masses with high fertilization rates until around the 10th egg mass, at which point fertilization decreased significantly (Fig. 3C). While it is difficult to draw conclusions about these patterns based on the small sample size (n=2, n=4), fertilization rates for all individuals monitored eventually declined to zero. Unexpectedly, two individuals (8 wpf, Fig 2C; J6xA, Fig 4A) produced a single fertilized egg out of hundreds or thousands of eggs laid. No other animals in the same treatment groups laid fertilized eggs (Fig. 2, Fig 4). Self fertilization is unlikely based on anatomy in this and other nudibranchs (Hadfield and Switzer-Dunlap 1984). More likely, low rates of sperm transfer, or insufficient development of the seminal receptacle (Fig. 5D) to maintain sperm contributed to this rare occurrence. Sperm present in these animals may also have degraded over the 7-8 week period between isolation and laying. However, other individuals in our experiments are able to store sperm for months (below).

Precocious sperm exchange requires that individuals be able to store sperm for some period of time. Sperm storage duration varies widely across the animal kingdom, from a matter of hours to years (see Orr and Brennan 2015 for review). Across the 15 individuals tracked in our study, we observed storage times of several months (Fig. 3C). In the nudibranch *Aeolidiella glauca*, isolated individuals were able to lay fertilized egg masses for 5-6 weeks (Karlsson and Haase 2002), and controlled matings of laboratory-reared *Aplysia californica* showed that individuals were able to produce fertilized egg masses from a single mating bout for up to 41 days (Ludwig and Walsh 2004). Most studies do not track animals for long enough to estimate the maximum duration of sperm storage; therefore, the number of fertilized egg masses produced by an individual might be a useful proxy. For example, in the nudibranch *Hermissenda crassicornis*, a single mating event provided enough sperm to lay about 2-3 fertilized egg masses (Rutowski 1983), and a single mating event in *A. california* produced ~9 fertilized egg masses (Ludwig and Walsh 2004). Rutowski used a cutoff of 50% fertilized eggs within a single egg mass to determine whether or not an egg mass is fertilized. Converting our data to match this cutoff, a week of mating in *B. stephanieae* was sufficient for an average of 14 fertilized egg masses. Obviously, more mating opportunities were available to individuals over the course of our experiment, though we did not quantify how many times animals mated. The ability to store sperm prior to the production of fertilized eggs and reciprocal mating opportunities produces high fecundity in individuals of *B. stephaniae*, similar to reports from in other heterobranchs.

Generally speaking, long term storage of sperm requires some physical adaptation (specific storage organ or otherwise), and may also include mechanisms for supporting and maintaining sperm (Orr and Brennan 2015). The reproductive system of nudibranchs is notoriously convoluted and complex, and in many species the reproductive tract includes areas where sperm from other individuals (allosperm) are stored (Hadfield and Switzer-Dunlap 1984; Wägele and Willan 2000). Here, we see distinct areas of sperm with heads directed towards the wall of the reproductive tract, identifying the seminal receptacles (Fig. 5G-I). No obvious changes are seen in the epithelium of the seminal receptacles between 6 mm and 9 mm juveniles (Fig. 5), but the appearance of sperm in both the seminal receptacles (allosperm) and ampulla (autosperm) in later-stage (9 mm and larger) animals indicates sperm is both given and received.

### Fertilization rate variation across individuals

Fertilization rates were variable within and across treatments, suggesting high rates of inter-individual variation in reproductive success. As previously noted, growth rates are highly variable in *B. stephaniae*, and depend on environmental conditions (ie. temperature, food availability, competition with conspecifics). This makes direct comparisons between age- and size-based experiments difficult, as well as suggesting a source of inter-individual variation. Overall, animals in the first (age-based) experiments had much lower rates of fertilization compared to individuals in the second (size-based) experiment, despite starting at about the same age as the large (9 mm) juveniles. These differences in fertilization rates are likely due to longer periods of sperm storage (2-4 weeks) in the age-based experiment, though may also have been due to preferable conditions (ie. higher temperatures, greater food availability, no competition with conspecifics) in the size-based experiment. In all experiments, fertilization declined markedly as individuals aged. Most large (9 mm) juveniles consistently laid egg masses with very high fertilization rates while most adults laid egg masses with variable fertilization rates (Fig. 4B). These adults (~7 months old) were ~3 months older than the large juveniles (~4 months old) at first laying. While age does not appear to affect sperm quality as animals laid egg masses of similar fertilization rates regardless of their mate (Fig. 4B), variable fertilization rates may be due to senescence and a decrease in egg quality in older animals. Senescence after oviposition is common in nudibranchs with many having lifespans of less than 6 months (Folino 1993; Schlesinger et al. 2009; Wolf and Young 2012). In our hands, the lifespan of *B. stephaniae* is between 6-18 months in the lab (data not shown), and lifespan is likely to be shorter in the wild. High predation on nudibranchs (Harris 1975) selects for early reproductive output, which is highly correlated with early senescence (Luckinbill et al. 1984). More practically, this result suggests focusing mating schemes/experiments on young adults in laboratory culture to maximize availability of high quality reproductive material.

## Conclusions

Detailed studies of reproduction, including reproductive timing and histology, provide important information about life history, and allow us to draw inferences about the lability of sexual systems. Similar studies in other species will be necessary to understand how widespread developmental variation in simultaneously hermaphroditic species is, and address how such systems may have evolved. Additionally, better understanding of the reproductive system of *B. stephanieae* will in turn enhance our ability to introduce genetic material in the *Berghia* system. For example, powerful new techniques such as ReMOT Control, which introduce transgenic components into adult animals, targeting the ovaries directly (Chaverra-Rodriguez et al. 2018, 2020), will require detailed understanding of reproductive processes in order to be effective. The increasing availability of genetic tools opens new avenues for exploring the diversity of sexual systems. However, such future work requires the solid foundations provided by detailed anatomical and reproductive data, such as what we provide here.

## Acknowledgements

We thank Nicholas Holland for his advice and use of his histological equipment. We also thank all members of the Lyons lab and the <u>*Berghia* BRAIN Project</u> team for their feedback, especially the Katz lab and Dr. Cheyenne Tait for the inspiration for the mixed-age mating experiments.

## Author Contributions

All authors contributed to the design of these experiments. NFT performed the nudibranch isolation experiments while isolating at home during the Covid-19 pandemic, and analyzed the data. NFT and JAG performed the histological sectioning. NFT and MPL prepared the initial draft of the manuscript. All authors contributed to and approved the final manuscript.

